# Host-directed FDA-approved drugs with antiviral activity against SARS-CoV-2 identified by hierarchical *in silico/in vitro* screening methods

**DOI:** 10.1101/2020.11.26.399436

**Authors:** Tiziana Ginex, Urtzi Garaigorta, David Ramírez, Victoria Castro, Vanesa Nozal, Ines Maestro, Javier García-Cárceles, Nuria E. Campillo, Ana Martinez, Pablo Gastaminza, Carmen Gil

## Abstract

The unprecedent situation generated by the COVID-19 global emergency prompted us to actively work to fight against this pandemic by searching for repurposable agents among FDA approved drugs to shed light into immediate opportunities for the treatment of COVID-19 patients.

In the attempt to proceed toward a proper rationalization of the search for new antivirals among approved drugs, we carried out a hierarchical *in silico/in vitro* protocol which successfully combines virtual and biological screening to speed up the identification of host-directed therapies against COVID-19 in an effective way.

To this end a multi-target virtual screening approach focused on host-based targets related to viral entry followed by the experimental evaluation of the antiviral activity of selected compounds has been carried out. As a result, five different potentially repurposable drugs interfering with viral entry, cepharantine, clofazimine, metergoline, imatinib and efloxate, have been identified.

## INTRODUCTION

Together with severe acute respiratory syndrome coronavirus (SARS-CoV) and Middle East respiratory syndrome coronavirus (MERS-CoV), SARS-CoV-2 is the third pathogenic and transmissible coronavirus emerged in humans. This new coronavirus (CoV) is the causative agent of the present pandemic of coronavirus disease named COVID-19 first reported in Wuhan (China) [1]. Since there is no effective treatment available and given the urgency of the pandemic, the repurposing of approved drugs is the only alternative to find a cure for the current emergency. In fact, several clinical trials are currently ongoing to prove the efficacy of old drugs in COVID-19 patients [2]. Such is the case of the drugs including in the SOLIDARITY clinical trial (remdesivir, hydroxychloroquine, lopinavir/ritonavir and interferon-beta1a), launched by the WHO in dozens of countries that showed little or no effects on hospitalized COVID-19 patients at proposed dose regimens[3]. Moreover, the only drug approved by the FDA for the treatment of extreme-ill patients is remdesivir [4], an antiviral originally developed for Ebola virus infection [5].

Although in principle not very innovative, drug repurposing is a promising approach to accelerate the drug discovery process which allows to increase the productivity of the pharmaceutical companies [6], and fill the gap existing in unmet diseases such as rare or infectious diseases [7–8]. In viral infections lacking of an effective treatment, drug repurposing combined with drug validation in animal models has enhanced the number of potential antivirals with known mechanism of action [9].

The COVID-19 global emergency has generated an unprecedent situation, which prompted scientists all around the world to actively work in all imaginable aspects related to SARS-CoV-2. In only few months, the knowledge of SARS-CoV-2 significantly increased and the available information today is quite large. Together with the efforts to better understand the epidemiology, virus structure and life cycle, several therapeutic targets to guide the drug discovery research have also emerged [10]. In this regard, it is remarkable the number of drug repurposing efforts trying to shed light into the COVID-19 patients treatment [11–12]. Today, far from initial opportunistic and mainly serendipitous discoveries in the drug repurposing field, a number of candidates have been proposed to be repurposed for COVID-19 based on different *in silico* and *in vitro* studies [13].

In the attempt to proceed toward a proper rationalization of the search for new antivirals among approved drugs, we here provide a hierarchical *in silico/in vitro* protocol, which successfully combines virtual and biological screening to speed up the identification of anti-SARS-CoV-2 agents in an effective way.

Moreover, as viral mutations represent one of the main challenges to overcome with antiviral therapies, we carried out a multi-target virtual screening protocol focused on druggable targets related to viral entry followed by biological screening against SARS-CoV-2 to identify host-directed therapies against COVID-19. In this regard, eight proteins mainly involved in SARS-CoV-2 entry and trafficking were considered.

Spike glycoprotein represents a crucial factor for virus entry and thus for virus tropism, virulence and pathogenesis [14–15]. For both SARS-CoV and SARS-CoV-2, cell-virus membrane fusion is promoted by the recognition of specific host proteins, or cell-binding agents such as the angiotensin-converting enzyme 2 (ACE2), which binds the receptor binding domain (RBD) located at the S1 subunit of the head region of the protein [15]. S priming is essential to promote membrane fusion. This process is catalyzed by specific host soluble proteases as the transmembrane serine protease 2 (TMPRSS2) mainly expressed in the surface of the airway epithelial cells [16]. TMPRSS2 was demonstrated to also cleave ACE2 [17–18], enhancing viral infectivity. Proteolytic cleavage of S is also promoted by other host proteases such as furin, which have cumulative effects of TMPRSS2-mediated S priming and SARS-CoV-2 entry [19–20]. Cathepsin L, a lysosomal cysteine protease of the papain family, is also involved in SARS-CoV and SARS-CoV-2 S priming. *In vitro* studies demonstrated that Cathepsin L can also perform proteolytic activity on the Spike glycoprotein when the other host proteases mainly involved in S priming are absent [21]. All these findings highlight the pivotal role exerted by host proteases in viral infection [22–23] thus confirming their inhibition as a valuable strategy to tackle COVID-19.

Furthermore, the adaptor-associated kinase 1 (AAK-1) and the cyclin G-associated kinase (GAK), members of the numb-associated kinase family (NAK), represent other two interesting drug targets against SARS-CoV-2 [24–25]. The main endosomal phosphatidylinositol-3-phosphate/phosphatidylinositol 5-kinase (PIKfyve) was also proposed to be related with intracellular trafficking of Ebola and SARS-CoV-2 viral particles [26].

Finally, the type 2 endo-lysosomal two-pore channel (TPC2), mainly expressed in late endosomes/lysosomes, mediates intracellular trafficking of coronavirus through the endo-lysosomal system. Accordingly, activation of TPC2 induces a calcium-dependent depolarization of the endo-lysosomal membrane, which is supposed to enhance S-driven membrane fusion [27]. In this context, TPC2 inhibitors such as verapamil [28], would be able to negatively affect depolarization thus reducing the fusogenic propensity during virus-host membrane fusion.

Considering this background, the US Drug Collection of 1789 compounds of FDA-approved drugs was then virtually screened towards the eight above mentioned targets and a total of 173 FDA repurposable drugs were selected from virtual screening and subsequently experimentally evaluated. Selection of these targets was motivated by their relevant role in virus life cycle, especially in virus recognition, entry and trafficking. Primary hits were validated using viral antigen detection in infected cells. Confirmed candidates were subsequently tested for their ability to interfere selectively with viral entry in a surrogate model of infection. This process led to the identification of cepharantine, clofazimine, metergoline, imatinib and efloxate as selective SARS-CoV-2 entry inhibitors, together with a panel of non-selective entry inhibitors that could be considered also for drug repurposing to treat COVID-19.

## RESULTS

### Virtual screening against selected targets

A hierarchical virtual screening (VS) approach was applied on crucial SARS-CoV-2 protein targets in the attempt to find repurposable agents from the original list of FDA approved drugs. Among all the proposed druggable targets for SARS-CoV-2, eight proteins responsible for virus entry and trafficking were selected in this study. A schematic representation of their role in virus entry and trafficking is displayed in **Figure 1**.

**Figure 1.**
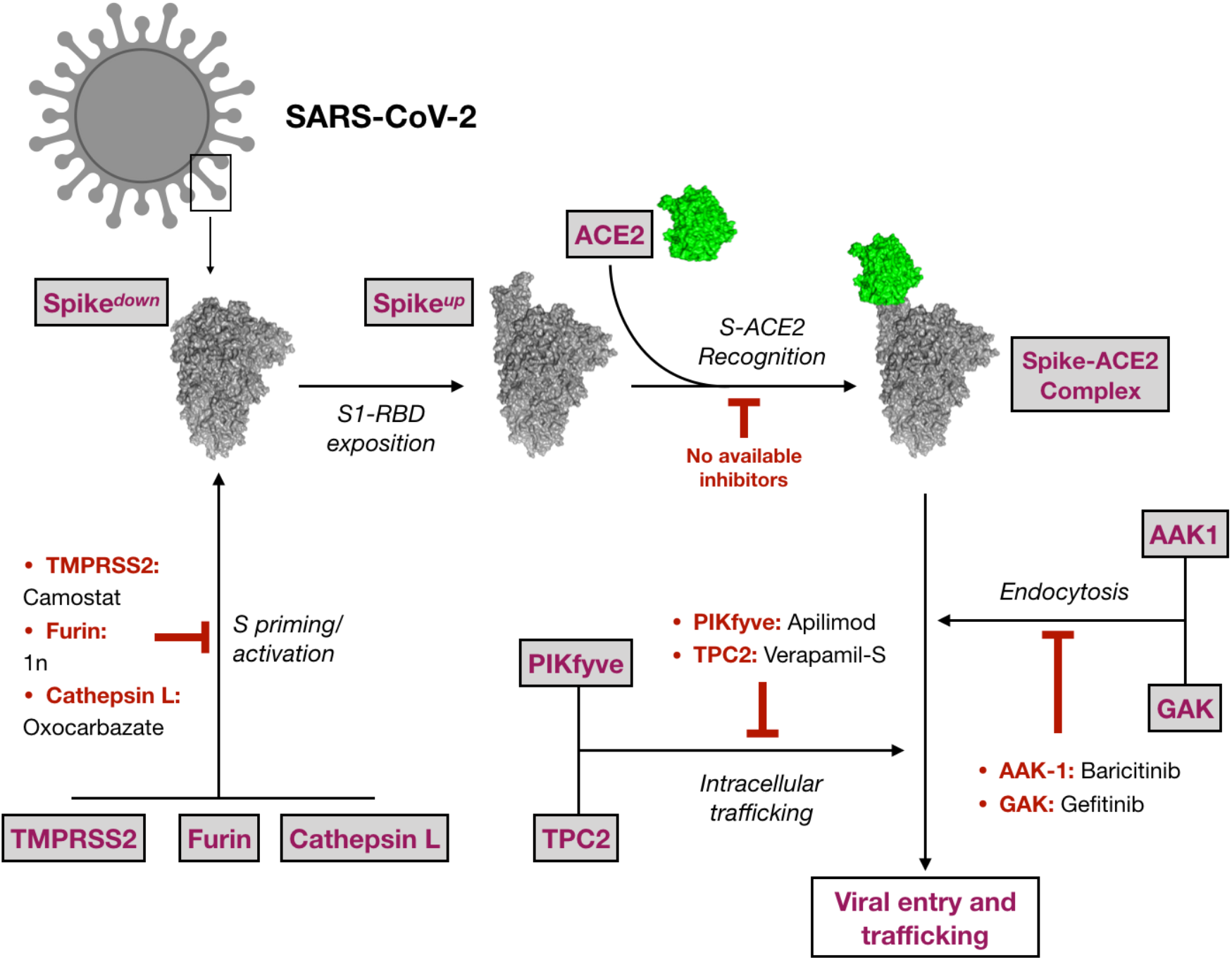
Schematic representation of the eight targets selected in this study and their role in virus entry. Representative inhibitors are also cited, when available.

A list of 1789 FDA-approved drugs was screened on all the previously cited targets (see Materials and Methods section for the computational details) to find effective antiviral compound candidates acting on SARS-CoV-2. Details about all the available PDB structures, druggable sites explored during VS and known inhibitors are reported in **Table S1** of the supporting information. The computational protocol applied in this study is shown in **Figure 2**.

**Figure 2.**
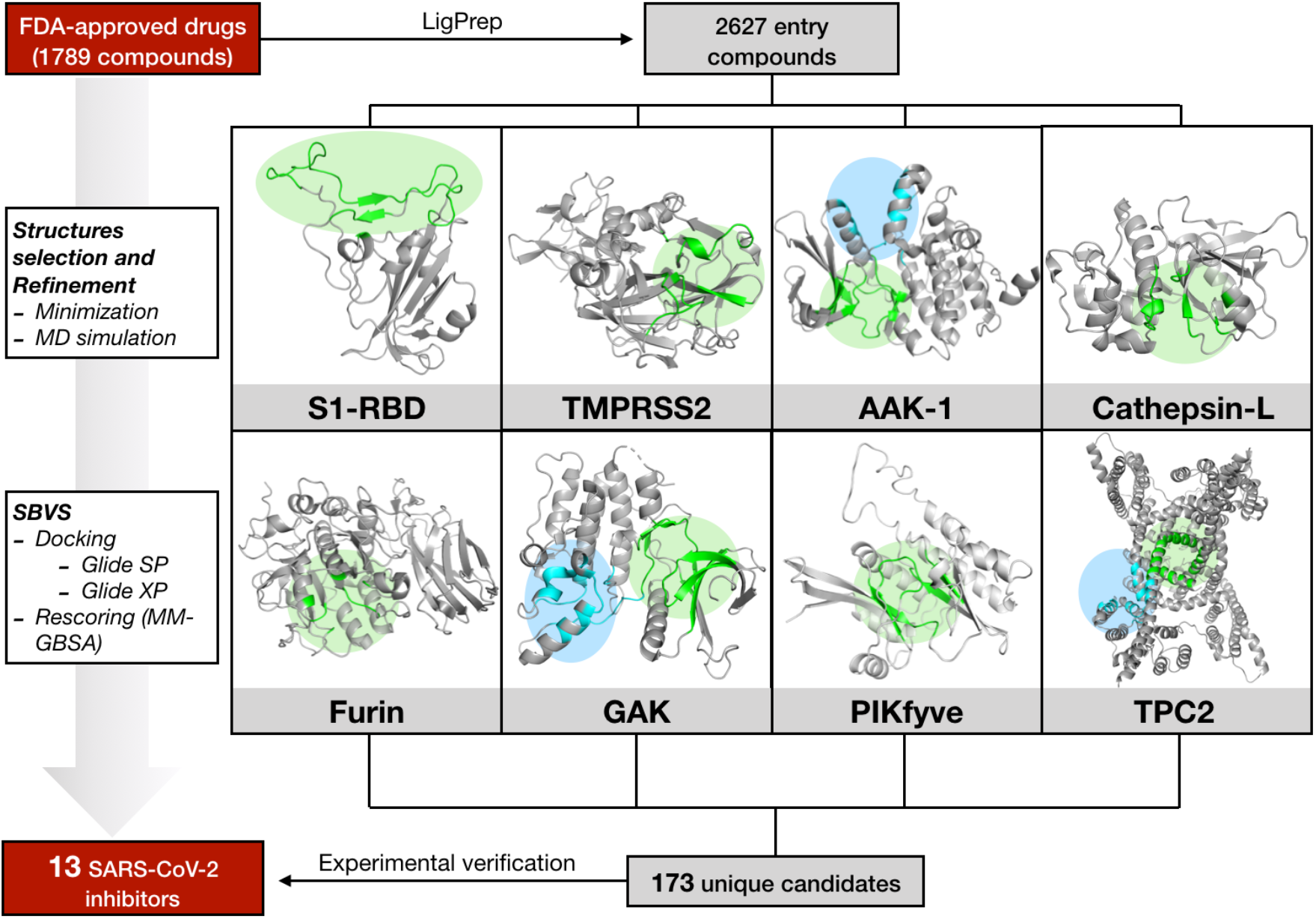
Schematic representation of the computational protocol applied in this study. For each target, green and blue circles respectively mark the active and the allosteric/secondary binding sites.

According to the scheme reported in **Figure 2**, all the systems were subjected to structural refinement by mean of energy minimization and the minimized structures were then used for VS. A special refinement was reserved to the S1-RBD domain of Spike and to the homology modelled structured of PIKfyve and TMPRSS2.

For PIKfyve enzyme, minimization of the homology modelled protein was realized in the presence of ATP substrate. This allowed to correctly reorient side chains for residues pertaining to the ATP binding site, preserving the geometry and shape of the cavity.

In case of S1-RBD and TMPRSS2, a further treatment based on molecular dynamic (MD) simulation in the NVT ensemble was applied. This allowed to properly explore local conformational flexibility of the ACE2 binding domain of S1-RBD and to refine the homology modelled structure of TMPRSS2 prior to virtual screening.

Trajectory analysis for S1-RBD and TMPRSS2 revealed a good stability along the MD simulation (**Figure S1** of the supporting information). For S1-RBD, a close analysis of the residues constituting the ACE2 recognition motif on the receptor binding region, revealed a mayor degree of fluctuation at the loop containing F154, N155 and Y157 (in dark blue in **Figure S1A**). Less mobility, generally lower than 1 Å, was observed in the other regions. For TMPRSS2 (**Figure S1B**), significant fluctuations were observed around the catalytic residues, H296, D345 and S441 (especially for loops in light blue, orange and green), which would be ascribable to the significant solvent exposition of the active site. For these two targets, the minimized structure and the most representative clusters (**Table S1** and **Table S2** of the supporting information) obtained from MD simulations were thus used for multi-conformation VS.

Accordingly, a total of 6 conformations for S-RBD of the Spike glycoprotein and 4 conformations for TMPRSS2 were considered for the following virtual screening (see Material and Methods section for additional information about clusters selection). For all the other targets, only the energy minimized crystallographic structure was considered (**Table S1** and **Table S2** of the supporting information).

All targets were subjected to a three-staged virtual screening protocol consisting on a preliminary docking by using the SP Glide docking algorithm, a second docking by applying the XP Glide docking algorithms and a final rescoring by applying the Prime MM-GBSA method. For each screened target, the 50 best ranked FDA drugs according to the MM-GBSA score were preliminary selected. Among them, compounds intended for a veterinary and/or cosmetic use, biocides, laxative or topical-administered drugs were not considered for SARS-CoV-2 *in vitro* assays. The complete list for the 173 selected drugs and their potential target(s) emerged from VS is shown in **Table S3** of the supporting information. These compounds were experimentally assayed for their SARS-CoV-2 antiviral potential within the framework of a *host-directed* COVID-19 antiviral therapy.

### SARS-CoV-2 antiviral candidate biological evaluation: experimental screening and prioritization

Selected candidates were evaluated for their antiviral activity in a cell culture model of SARS-CoV-2 infection. Cytopathic effect (CPE) was determined in Vero-E6 cells, which are particularly susceptible to SARS-CoV-2 infection with a high viral load resulting in general cell death after 72 hours of infection. Cell death can be readily delayed and even prevented by treatment with reference antiviral compounds and may be used to identify new antivirals [12]. Thus, antiviral activity of new drugs can be revealed by the ability of a given compound to protect the cell monolayer upon infection. To effectively quantify the antiviral potential of FDA-approved drugs identified by the multi-target virtual screening described above, we tested the 173 candidates for their ability to protect Vero-E6 cells from virus-induced cell death at a fixed concentration of 10 μM. Infected cell monolayer integrity was assessed by crystal violet staining 72 hours after inoculation at a multiplicity of infection (MOI) of 0.001. This analysis revealed 26 compounds that prevented virus-induced cell death at 10 μM and 7 compounds that were cytotoxic at this concentration (**Table S3** of the supporting information).

Both sets of compounds were counterscreened in a dose-response experiment to determine the range of concentrations capable of protecting the cell monolayer and to confirm their antiviral potential. Only one of the cytotoxic compounds, lanatoside C, revealed antiviral activity at lower concentrations, while the other 6 cytotoxic drugs did not reveal any protective activity. Six of the primary hits (posaconazole, thiostrepton, dipyridamole, hycanthone, gefitinib and pirenpirone) could not be unequivocally confirmed as they did not confer full protection against virus-induced cytopathic effect at any of the assayed doses. Moreover, eight of the candidates (loratadine, ivermectin, terfenadine, lapatinib, carvedilol, tilorone, reserpine and amoxapine) showed a narrow therapeutic window, since they conferred protection at a unique dose. Thus, these compounds were not further characterized. Niclosamide, digoxin, penfluridol, clofazimine, cepharantine, imatinib, pimozide, metergoline, mycophenolate mofetil, lanatoside C, efloxate, ebastine and protoporphyrin IX clearly prevented SARS-CoV-2-induced cytopathic effect at more than one dose and were selected for further characterization.

### Antiviral activity of selected candidates

The antiviral candidates have been selected based on their ability to prevent virus-induced cell death, which is an indirect assessment of virus infection efficiency. In order to directly confirm the antiviral activity of the selected compounds, viral antigen expression was assessed in the presence of the candidate compounds by immunofluorescence microscopy in SARS-CoV-2 infected cells using an antibody raised against SARS-CoV-2 nucleoprotein (N). Infections in the presence of a range of compound concentrations were carried out at a MOI of 0.01 and cells were fixed at 24 hours post inoculation, time at which no virus-induced cytopathic effect is observed. At this time of infection and MOI, SARS-CoV-2 infection has locally spread in Vero-E6 and infection efficiency may be estimated by the expression of N protein. Staining of cell nuclei using DAPI (4,6-diamidino-2-phenylindole) allows evaluation of the cell number to verify that antiviral activity occurs at non-cytotoxic concentrations. Dose response datasets (**Figure S2**) were used to calculate EC_50_ and EC_90_ values, corresponding to the concentration of compound that causes 50% or 90% reduction of viral antigen accumulation respectively (Table 1). Remdesivir, a broad-spectrum nucleotide analog with anti-SARS-CoV-2 activity *in vitro* and *in vivo* was used as control and the estimated EC_50_ (1.6 μM) was similar to that previously reported in this cell line [29], indicating that the method is appropriate to estimate the potency of the compounds.

**Table 1.**
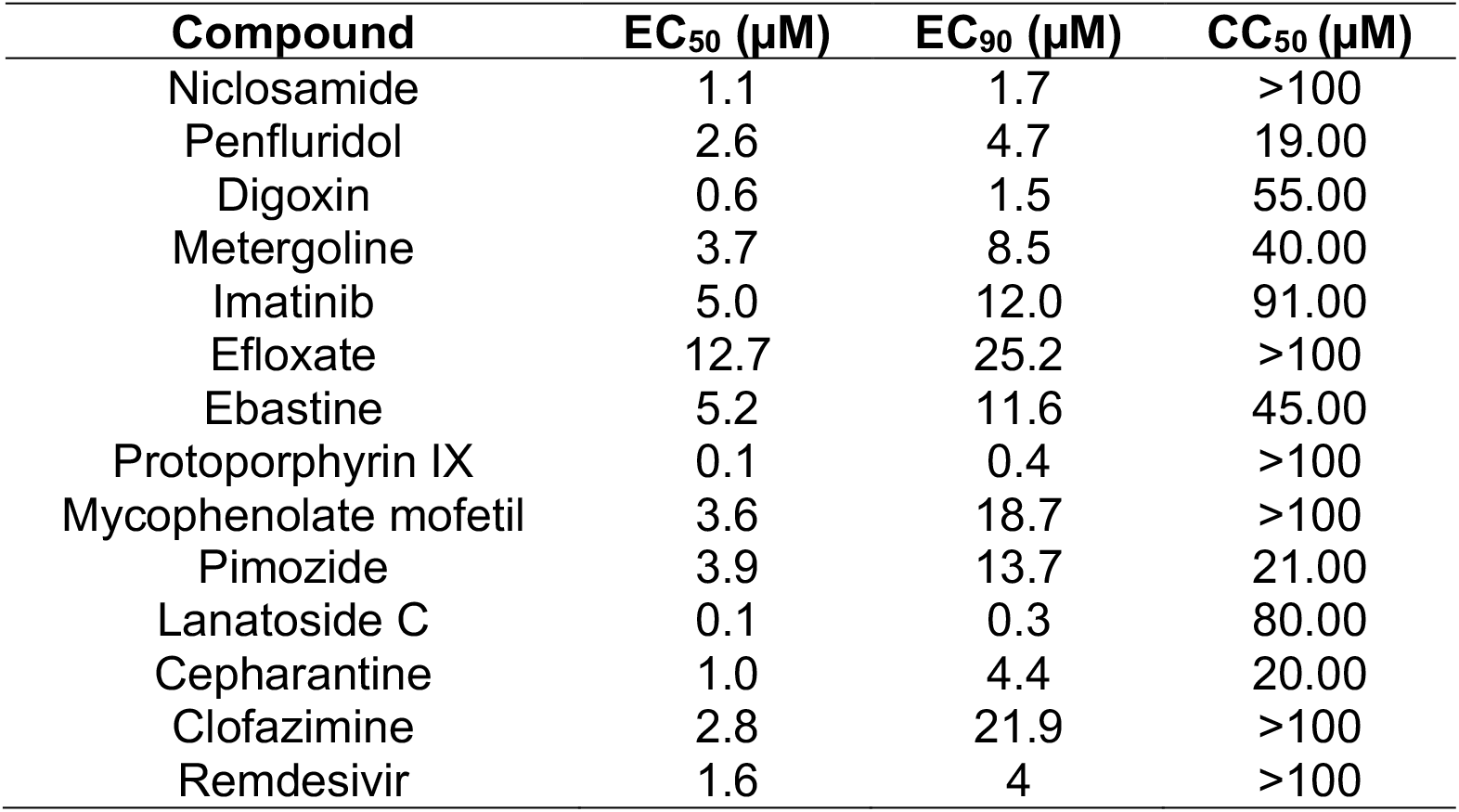
Potency and cytotoxicity indexes of the confirmed primary hits. Potency (EC_50_; EC_90_) and cytotoxicity (CC_50_) were calculated from dose-response experiments in which infection efficiency was determined by viral antigen accumulation and MTT activity respectively (see Figures S2 and S3).

To determine the impact of the antiviral candidates on overall cell viability, proliferation and cytotoxicity, we performed an MTT assay with a wide range of compound concentrations to estimate the CC_50_ value or the compound concentration that causes the 50 % of cytotoxicity. **Figure S3** shows the results of the MTT assay and the inferred CC_50_ values are shown in Table 1. Based on these indexes, substantial antiviral activity was observed for all the compounds at non-cytotoxic doses (**Table 1**). However, as it can be observed in **Figure S3**, treatment of the cells with niclosamide, clofazimine or protoporphyrin IX resulted in a marked, dose-dependent elevation of the MTT activity at a broad range of concentrations, suggesting that they either interfere with the assay itself or that they cause substantial stress to the cells without killing them. In the case of protoporphyrin IX and clofazimine, both compounds are colored and stain the cells significantly at the highest concentrations. While this may interfere with the interpretation of the MTT results at the highest doses, these compounds display antiviral activity at lower doses, thus reducing the concern for this phenomenon. In contrast, the lowest dose of niclosamide with antiviral activity shows substantial MTT elevation (**Figure S2** and **S3**), probably owing to its reported mitochondrial uncoupling [30] and oxidative stress induction capacity [31].

Furthermore, elevation of the MTT activity at doses preceding marked cytotoxicity was observed in imatinib, penfluridol, pimozide, efloxate, ebastine and metergoline, probably as a transient adaptation to compound-induced stress [32]. Anyhow, and with the exception of pimozide and niclosamide, all the above-mentioned drugs showed clear antiviral activity (EC_90_) associated with cells displaying normal MTT activity (**Figure S3**).

### Evaluation of anti-SARS-CoV-2 drugs as entry inhibitors

As the SARS-CoV-2 inhibitors described above were identified by a multi-target host-based entry targets screening, their ability to interfere with SARS-CoV-2 entry was evaluated in a surrogate model of infection based on retroviral vectors pseudotyped with SARS-CoV-2 Spike envelope glycoprotein. This system encompasses the production of reporter retroviral vectors pseudotyped with the envelope glycoprotein S (Spp), which is a major determinant of SARS-CoV-2 entry, mediating receptor recognition, internalization and viral membrane fusion. This system enables evaluation of virus entry efficiency as a function of the reporter gene activity (luciferase), which is strictly dependent on the presence of a functional viral glycoprotein.

Entry efficiency was evaluated in the presence of the EC_90_ of the candidates, except for pimozide, which was evaluated at 6.25 μM given the elevated MTT activity observed at the EC_90_ (**Figure S3**). This analysis revealed that, as expected, the polymerase inhibitor remdesivir did not interfere with the entry process (**Figure 3A**). Similarly, niclosamide, digoxin, lanatoside C, penfluridol and pimozide did not show any marked reduction in virus entry efficiency (**Figure 3A**) at doses capable of reducing infection efficiency by one order of magnitude (**Table 1**). Thus, these results suggest that these compounds interfere with virus infection by a mechanism that does not clearly interfere with Spike-mediated entry at the assayed doses. In contrast, a significant reduction in Spp entry was observed in the presence of protoporphyrin IX, cepharantine, efloxate, clofazimine, metergoline, imatinib, mycophenolate mofetil and, although modestly, ebastine (**Figure 3B**). To determine if the observed inhibition is selective for SARS-CoV-2 Spike-mediated entry, pseudotypes based on vesicular stomatitis virus (VSV-G) or RD114 glycoprotein were studied in parallel. VSV-G pseudotypes use the endocytic pathway to enter the cells, although using a different receptor that S-pseudotypes and with a remarkable efficiency in many cell types [33]. On the other hand, RD114-pseudotypes, are internalized after direct fusion of the viral envelope with the cell plasma membrane and do not follow the endocytic route [34]. Mycophenolate mofetil interfered with entry of all three tested retroviral pseudotypes, probably owing to its demonstrated ability to inhibit DNA and RNA viral infections by depleting cell nucleotide pools, a function that may interfere with this retrovirus-based assay [35]. Protoporphyrin IX interfered similarly with Spp and VSVpp and to a lesser extent with RD114pp entry, suggesting non-selective interference with virus enveloped internalization, in agreement with previously reported data [36]. Partial selectivity was observed for cepharantine, efloxate, clofazimine, metergoline, imatinib and ebastine (**Figure 3B**).

**Figure 3.**
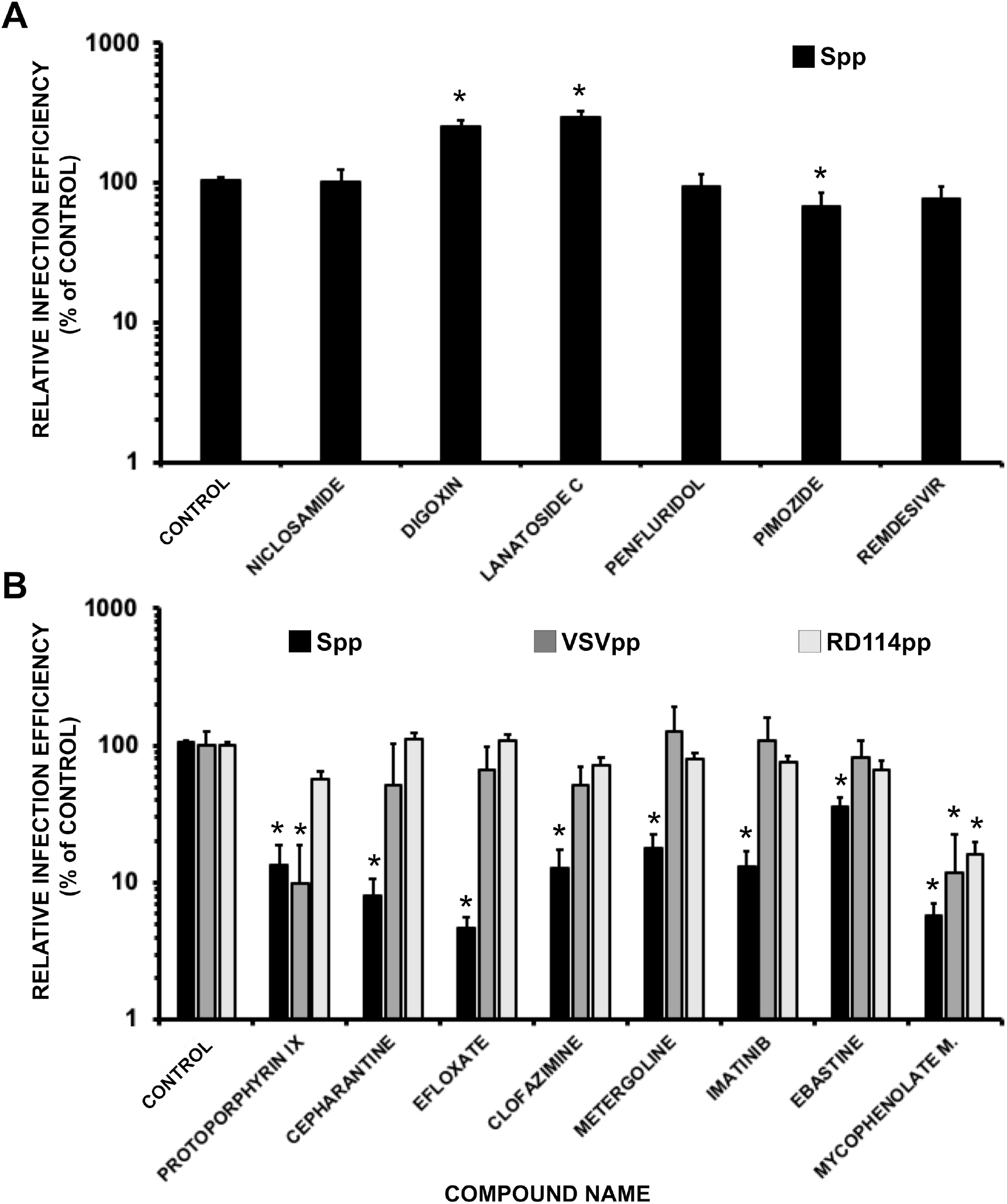
Antiviral candidates interfere with viral entry of SARS-CoV-2 pseudotypes. A) Retroviral vectors pseudotyped with SARS-CoV-2 Spike glycoprotein were used to inoculate Vero-E6 cells in the presence of the candidates at the EC_90_ (see Table 1). Forty-eight hours later, total cell lysates were assayed to determine luciferase activity as a reporter activity for viral entry. B) Compound selectivity was assayed also using VSVpp and RD114pp for the compounds interfering with Spp. Data are shown as average and SEM of a minimum of four biological replicates (N=4). Statistical significance was evaluated using a one-way ANOVA and a Dunnet’s post-hoc test.

Overall, our data suggest that interference with viral entry is a major contributor to the antiviral activity shown by cepharantine, efloxate, clofazimine, metergoline and imatinib and against SARS-CoV-2 in cell culture.

## DISCUSSION

Since its first detection in December 2019 in Wuhan, the capital of China’s Hubei province, COVID-19 has spread worldwide rapidly. The outbreak was declared a Public Health Emergency by WHO on 30 January 2020 and since then, utmost efforts were made by the international scientific community in the attempt to find an effective cure. The full characterization of the SARS-CoV-2 viral genome by Fuk-Woo Chan J. and collaborators [37], followed by crystallization of most of its viral components offered the structural bases to search for an effective treatment.

Vaccines represent the gold standard long-term choice to fight SARS-CoV-2 pandemic and COVID-19. However, vaccination campaigns will require a coordinated effort worldwide and full protection of the general population may be delayed for years or never be reached. Moreover, the emergence of new virus variants may reduce vaccine efficacy. Thus, pharmacological treatment of the infections with small molecules is a valid approach, but it is affected by important disadvantages, such as low potency and emergence of drug-resistant virus variants, especially when applied as monotherapy. These limitations could be dampened by the application of broad-spectrum antiviral agents simultaneously acting on more than one target at the same time [38]. Furthermore, to reduce the likelihood of resistance in future treatments, the design of antivirals able to block host targets involved in viral infection is an emerging and promising strategy [39]. In fact, this is the approach followed in this study. Thus, we screened *in silico* the same chemical library against eight different entry SARS-CoV-2 targets, being all of them human proteins.

The US Drug Collection of 1789 compounds of FDA-approved drugs was screened toward these targets, which consisted on the proteases TMPRSS2, Furin and Cathepsin L, the kinases AAK1, GAK and PIKfyve as well as the two-pore ion channel TPC2. Additionally, the receptor binding domain (S-RBD) of the viral Spike (S) glycoprotein, which is recognized by the host protein, ACE2 during virus attachment, was included in the analysis.

Following this trend, a hierarchical host-directed virtual screening protocol was applied to select potential anti-SARS-CoV-2 drugs based on the above-mentioned host targets with the aim to find a host-based therapy for COVID-19 capable to interfere with virus attachment, endocytosis and trafficking. In this regard, 173 FDA approved drugs were selected from the multi-target *in silico* virtual screening and finally tested against SARS-CoV-2 viral infection using a high-throughput screening (HTS) protocol which was optimized for this work.

The potential antiviral activity of the selected FDA-approved drugs selected during VS was first evaluated in a cell culture model of SARS-CoV-2 infection at a fixed concentration of 10 μM. Vero-E6 cells were selected because of the proven susceptibility to the infection by this coronavirus. This preliminary assay yielded 26 hits (**Table S3**) and subsequent dose-response experiment to determine the range of protective concentration allowed confirmation of 13 candidates with antiviral activity at non-cytotoxic doses for further studies (**Table 1**).

The aim of this study was to determine if any of the antiviral candidates interferes with viral entry, as predicted by the bioinformatic analysis. A specific assay consisting on S protein pseudotyped retroviral vectors was set up to gain deeper knowledge about the potential ability of identified antivirals to inhibit SARS-CoV-2 entry. In these experiments, we confirmed that cepharantine, imatinib, efloxate, clofazimine, and metergoline prevent viral infection primarily by interfering with viral entry (**Figure 3**).

Cepharantine was approved in Japan to treat alopecia [40], which was proposed in the last months as anti-COVID-19 therapy based on theoretical and *in vitro* results [11]. Our *in silico* results showed that cepharantine could be a potential inhibitor of furin and TPC2, being its biological action involved not only in first entry phases but also in the escape from late endosomes, a mechanism that is compatible with the results obtained in the surrogate model of viral entry here presented (**Figure 3B**) and a recently published report [41].

Due to the involvement of Abl pathway in viral infections, imatinib was proposed as anti-SARS-CoV-2 and clinical trials were started since the first moment of the pandemic [42], although no experimental evidence of antiviral activity was reported. At the time of writing this manuscript, *in vitro* activity against SARS-CoV-2 has been described [43] in agreement with the results here presented.

To our knowledge, no antiviral activity has ever been reported for the vasodilator efloxate. Our virtual screening shows that this drug could potentially inhibit AAK1 and GAK kinases involved in early endosome entry. Similarly, no antiviral activity has been reported for metergoline against SARS-CoV-2, which our VS predicts to interfere with host proteases involved in viral entry.

Clofazimine, used as an antimicrobial agent, also showed consistent antiviral activity as previously reported in a similar infection system [12]. Our data suggest that clofazimine interferes selectively with SARS-CoV-2 (**Figure 3B**).

Five broad-spectrum antiviral compounds, niclosamide, digoxin, protoporphyrin IX, mycophenolate mofetil and lanatoside C were considered in this study to determine their impact on viral entry. We confirmed antiviral activity for all five compounds. However, we did not observe interference with SARS-CoV-2 entry for lanatoside C, niclosamide or digoxin at the EC_90_, suggesting that interference with viral entry is not a major contributor to infection inhibition at the assayed concentrations.

Consistent with our results digoxin has been recently shown to interfere with viral replication at post-entry stages [44]. Also, lanatoside C has previously been shown to display antiviral activity against different RNA viruses at post-entry steps [45]. Finally, niclosamide displays antiviral activity against a broad range of viruses, including SARS-CoV [46] and it has been proposed that it could interfere with viral entry by preventing endosomal acidification. We failed to show this effect since we tested the compound at doses devoid of any alteration of the MTT values.

Owing to their previously described broad antiviral activity, protoporphyrin IX [36] and mycophenolate mofetil [47] displayed relatively non-specific inhibition of the retroviral pseudotyped entry and precluded concluding on their genuine impact on SARS-CoV-2 viral entry.

## CONCLUSIONS

Overall, the use of an FDA-approved chemical library allowed us to check the robustness and reproducibility of our protocol, a multi-target virtual screening following by a solid experimental cascade of biological assays. Our study allowed the identification and experimental validation of valuable candidates to be repurposed as potential COVID-19 therapy such as cepharantine, efloxate, metergoline, imatinib, clofazimine, digoxin, protoporphyrin IX, and lanatoside C. Moreover, a potential mechanism of action for these drugs was also proposed by *in silico* VS analyses as they would be able to modulate some of the host proteins involved in the entry process of SARS-CoV-2 and was experimentally supported for cepharantine, imatinib, efloxate, clofazimine, and metergoline. In summary, we have identified a list of five drugs ready to be validated in clinical trials as SARS-CoV-2 infection inhibitors. In case of positive results from clinical trials with COVID-19 patients, these compounds may promote a new era of antiviral agents potentially able to combat the current COVID-19 pandemic, but also future outbreaks of high pathogenic viruses, which would share a common entry pathway as infection mechanism.

## MATERIALS AND METHODS

### Computational studies

#### The drug-dataset

A starting list of 1789 FDA-approved drugs (US Drug Collection, MicroSource Discovery Systems) has been prepared for VS with the LigPrep and Epik modules of Maestro suite [48]. Accordingly, all possible ionization states at pH 7.2 ± 2.0 have been predicted for each compound. Original chirality has been retained. This led to a total of 2627 compounds. The force field, OPLS3 [49] has been used to define all the generated compounds.

#### Protein targets and MD simulations

The X-ray crystal structure of SARS-CoV-2 spike receptor binding domain in complex with ACE2 (PDB ID: 6M0J) [50] has been used as model for the S1-RBD-ACE2 recognition surface.

For TMPRSS2, the homology modelled extracellular region of the protein was obtained from the Swiss-Model repository [51]. The model was obtained from the serine protease hepsin (PDB ID: 5CE1), which shares the 34% of sequence identity with the target protein, TMPRSS2.

Amber 18 [52] was used to explore the local conformational flexibility of the S1-RBD of Spike and to refine the homology modelled structure of TMPRSS2. The ff14SB force field [53] was used to define the proteins which were embedded in a truncated octahedral TIP3P [54] water box in a layer of 22 Å and neutralized by adding chlorine counterions. Disulphide bonds were built by using the “bond” command in tleap.

Protonation states for titratable residues were set according to Propka [55] predictions at pH 7.3. Systems were energy minimized in three steps involving firstly all hydrogen atoms, then water molecules, and finally all the system. For the final step, a maximum of 50,000 (5,000 iterations with steepest descent and the rest with conjugate gradient) were run. Thermalization of the minimized systems from 0 to 300 K was accomplished in five steps, the first being performed at constant volume and the rest at constant pressure. Langevin dynamics with a collision frequency of 1.0 ps^−1^ was applied for temperature regulation during thermalization. Prior to MD, 5 ns of equilibration at constant pressure were run to properly stabilize the systems. A total of 100 ns of MD production were generated in the NVT ensemble and in periodic boundary conditions for both systems. A time step of 2 fs was set for saving trajectories.

The SHAKE algorithm [56] was applied to constrain bonds involving hydrogen atoms. Cut-off for non-bonded interactions was set to 10 Å. Electrostatic interactions beyond the cut-off within the periodic box were computed by applying the Particle Mesh Ewald (PME) method [57]. The weak-coupling algorithm with a time constant of 10.0 ps was used to stabilize the temperature during the simulation. Trajectories analysis and clusterization were done by using the CPPTRAJ module of Amber18. For clustering analysis, a total of 10 clusters were preliminary searched by using the average linkage algorithm, which uses the average distance between members of two clusters [58]. Representative structure for each cluster was represented by the average structure. Cut-off for determining local density was set at 4 angstroms. Parameters for clusterization were adapted considering both trajectory and protein length.

For human PIKfyve, the structure of the protein has been obtained by homology modeling by using the crystal structure of zebrafish Phosphatidylinositol-4-phosphate 5-kinase alpha isoform with bound ATP/Ca^2+^ (PDEB code: 6CMW), which shares the 28% of global sequence identity [59]. To proper refine the ATP binding site in the homology modelled PIKfyve enzyme, ATP has been accommodated in its binding site by using the template complex as reference and the so derived PIKfyve-ATP complex has been then energy minimized. The ATP parameters for minimization with Amber18 were taken from the Amber parameter database of the Bryce group [60–61]. The complete list for all the crystallographic structures used during VS is reported in **Table S1**.

#### Structure-based Virtual Screening (SBVS) and MM-GBSA rescoring

The complete FDA database of 1789 approved drugs was screened against the previously described targets. Representative 3D-structures/clusters for S1-RBD and TMPRSS2 were selected from MD simulations. For cluster selection on S-RBD, clusters with a population higher than 5% (clusters 0-4) were considered for VS with the aim to enhance the exploration of the conformational variability of the receptor binding site of S protein. In case of TMPRSS2, only those clusters where the active site was in an open state were considered, 3 clusters (cluster 0-2) were finally selected for VS. For both systems, the minimized structure was also considered for VS.

For the rest of the screened systems (AAK-1, Cathepsin-L, furin, GAK, PIKfyve and TPC2), the minimized crystallographic structures were prepared for virtual screening with the protein preparation wizard from the Maestro suite, applying the OPLS3e force field [49] with default parameters. The grid boxes were centered on the active site for each target (see **Table S1**) using default parameters for receptor grid generation.

SBVS was then performed by using a pipeline which included 3 stages. The first one consisted in massive docking simulations employing the Glide software [62] and the Standard Precision (SP) method. In this first stage, an enhanced sampling approach was used, and 5 poses were generated per compound state. The best 50% of compounds (according to the scoring function) were kept and used for the second stage, where the Extra Precision (XP) method was employed. In the second stage, 25% of the best-ranked solutions were kept. Rescoring was performed in the third stage with Prime MM-GBSA method [63].

### SARS-CoV-2 infection assays

All infection experiments were performed by inoculating Vero-E6 cells seeded onto 96-well plates (2×10^4^ cells/well) with the SARS-CoV-2 strain NL/2020 (kindly provided by Dr. R. Molenkamp, Erasmus University Medical Center Rotterdam) at low multiplicity of infection (MOI) of 0.01 or 0.001, as indicated below. Cultures were maintained at 37 °C in a 5% CO_2_ incubator for different lengths of time depending on the experiment. Compounds were diluted from 10 mM stock solutions in complete media containing 2% FBS to achieve the indicated final concentrations.

#### Cell monolayer protection assays

For the primary screening, Vero-E6 cell monolayers were inoculated at MOI 0.001 in the presence of 10 μM of each compound in duplicate wells. Seventy-two hours later the cells were fixed and stained using crystal violet. Compounds that protect from the virus induced cell death were selected for further experiments. A wide range of two-fold dilutions of the compound (final concentration from 50 to 0.78 μM) were used in subsequent experiments to determine the maximum and minimum protective concentrations as indicated above. Only compounds conferring full protection at two consecutive dilutions (2-fold dilutions) were considered for further characterization.

#### Evaluation of the antiviral activity immunofluorescence microscopy

VeroE6 were seeded onto 96-well plates as described above and infected in the presence of the indicated compound dose (MOI 0.01). Twenty-four hours post infection, cells were fixed for 20 minutes at RT with a 4% formaldehyde solution in PBS, washed twice with PBS and incubated with incubation buffer (3% BSA; 0.3% Triton X100 in PBS) for 1 hour. A monoclonal antibody against N protein was diluted in incubation buffer (1:2000; Genetex HL344) and incubated with the cells for 1 hour, time after which the cells were washed with PBS and subsequently incubated with a 1:500 dilution of a goat anti-rabbit conjugated to Alexa 488 (Invitrogen-Carlsbad, CA). Nuclei were stained with DAPI (Life Technologies) during the secondary antibody incubation using the manufacturer’s recommendations. Cells were washed with PBS and imaged using an automated multimode reader (TECAN Spark Cyto; Austria).

Clofazimine interfered with the fluorescence-based assay due to its intrinsic fluorescence [64] at highest concentrations. Thus, this assay was performed using a colorimetric readout similar to what has previously been described [65], using a primary human monoclonal anti-S antibody (kindly provided by L. A. Fernández and J.M. Casasnovas (CNB-CSIC; Madrid) and a secondary goat anti-human Fc antibody conjugated with horseradish peroxidase.

EC_50_ and EC_90_ values were obtained using PROBIT regression method using IBM SPSS Software Package (version 26) using the average infection efficiency values from three biological replicates.

#### Cytotoxicity measurement by MTT assays

Vero-E6 cell monolayers were treated with a wide range of compound concentrations (from 50 to 0.78 μM) and forty-eight hours later they were subjected to MTT assays using standard procedures [66]. CC_50_ values were graphically interpolated from dose-response curves obtained with three biological replicates.

### SARS-CoV-2 Spike protein-pseudotyped retroviral vectors

Retroviral particle production pseudotyped with different viral envelopes has previously been described [67–68]. Packaging plasmids, vesicular stomatitis virus (VSV) G and RD114 glycoprotein expressing plasmids were kindly provided by Dr. F. L. Cosset (INSERM, Lyon). SARS-CoV-2 S expressing plasmid was obtained from Jose María Casasnovas and Juan García Arriaza (CNB-CSIC). Particles devoid of envelope glycoproteins were produced in parallel as controls.

For SARS-CoV-2 Spike pseudotyped particle (SARS2pp) entry experiments, Vero-E6 cells (10^4^ cells/well) were seeded onto 96-well plates the day before. Compounds were diluted in complete media [(DMEM supplemented with 10 mM HEPES, 1x non-essential amino acids (Gibco), 100 U/ mL penicillin-streptomycin (Gibco) and 10% Fetal Bovine Serum (heat-inactivated at 56 °C for 30 min)] to achieve a 2x concentration. Fifty microliters (50 μL) of the SARS2pp, VSVpp or RD114 retrovirus dilutions were mixed 1:1 with 50 μL of the 2x compound dilutions to achieve the EC_90_. One hundred μL of the mixture was applied onto the Vero E6 cell monolayer in at least four biological replicates and cells were cultured at 37 °C in a 5% CO_2_ incubator. Twenty-four hours later, cell culture media was replaced with compound-free media. Forty-eight hours post-inoculation, cells were lysed for luciferase activity determination using Luciferase Assay System (Promega) and a luminometer. Relative infection values were determined by normalizing the data to the average relative light units detected in the vehicle control cells.

## Acknowledgments

Funding from CSIC (201980E024, 202020E103 and PIE-RD-COVID-19 ref. E202020E079), EVA (European Virus Archive; grant agreement N° 871029), ANID (Agencia Nacional de Investigación y Desarrollo de Chile, grant N° COVID0199), and CONICYT-PCI (Comisión Nacional de Investigación Científica y Tecnológica de Chile – Programa de Cooperación Internacional, grant N° REDES190074) is acknowledged. Dr. R. Molenkamp (Erasmus University Medical Center, Rotterdam, Netherlands; participant of the EVA-GLOBAL project) is acknowledged for the SARS-CoV-2 strain NL/2020 virus. Dr. F. L. (Inserm-Lyon) and Dr. Casanovas (CNB-CSIC) are acknowledged for the materials required to produce retroviral pseudotypes. Dr. Fernández is acknowledged for providing anti-S antibodies. I. M. was funded by H2020-MSCA-ITN-2017 (grant no. 765912) and V. N. holds a pre-doctoral FPU grant (FPU16/04466).

## References

1. Zhou, P.; Yang, X. L.; Wang, X. G.; Hu, B.; Zhang, L.; Zhang, W.; Si, H. R.; Zhu, Y.; Li, B.; Huang, C. L.; Chen, H. D.; Chen, J.; Luo, Y.; Guo, H.; Jiang, R. D.; Liu, M. Q.; Chen, Y.; Shen, X. R.; Wang, X.; Zheng, X. S.; Zhao, K.; Chen, Q. J.; Deng, F.; Liu, L.; Yan, B.; Zhan, F. X.; Wang, Y. Y.; Xiao, G. F.; Shi, Z. L., A pneumonia outbreak associated with a new coronavirus of probable bat origin. Nature 2020, 579, 270–273.

2. Saul, S.; Einav, S., Old drugs for a new virus: Repurposed approaches for combating COVID-19. ACS Infect Dis 2020, 6, 2304–2318.

3. Pan, H.; Peto, R.; Henao-Restrepo, A. M.; Preziosi, M. P.; Sathiyamoorthy, V.; Abdool Karim, Q.; Alejandria, M. M.; Hernández García, C.; Kieny, M. P.; Malekzadeh, R.; Murthy, S.; Reddy, K. S.; Roses Periago, M.; Abi Hanna, P.; Ader, F.; Al-Bader, A. M.; Alhasawi, A.; Allum, E.; Alotaibi, A.; Alvarez-Moreno, C. A.; Appadoo, S.; Asiri, A.; Aukrust, P.; Barratt-Due, A.; Bellani, S.; Branca, M.; Cappel-Porter, H. B. C.; Cerrato, N.; Chow, T. S.; Como, N.; Eustace, J.; García, P. J.; Godbole, S.; Gotuzzo, E.; Griskevicius, L.; Hamra, R.; Hassan, M.; Hassany, M.; Hutton, D.; Irmansyah, I.; Jancoriene, L.; Kirwan, J.; Kumar, S.; Lennon, P.; Lopardo, G.; Lydon, P.; Magrini, N.; Maguire, T.; Manevska, S.; Manuel, O.; McGinty, S.; Medina, M. T.; Mesa Rubio, M. L.; Miranda-Montoya, M. C.; Nel, J.; Nunes, E. P.; Perola, M.; Portolés, A.; Rasmin, M. R.; Raza, A.; Rees, H.; Reges, P. P. S.; Rogers, C. A.; Salami, K.; Salvadori, M. I.; Sinani, N.; Sterne, J. A. C.; Stevanovikj, M.; Tacconelli, E.; Tikkinen, K. A. O.; Trelle, S.; Zaid, H.; Røttingen, J. A.; Swaminathan, S., Repurposed antiviral drugs for Covid-19 - interim WHO Solidarity trial results. N Engl J Med 2021, 384, 497–511.

4. https://www.fda.gov/news-events/press-announcements/coronavirus-covid-19-update-fda-issues-emergency-use-authorization-potential-covid-19-treatment (accessed March 3, 2021).

5. Mulangu, S.; Dodd, L. E.; Davey, R. T., Jr.; Tshiani Mbaya, O.; Proschan, M.; Mukadi, D.; Lusakibanza Manzo, M.; Nzolo, D.; Tshomba Oloma, A.; Ibanda, A.; Ali, R.; Coulibaly, S.; Levine, A. C.; Grais, R.; Diaz, J.; Lane, H. C.; Muyembe-Tamfum, J. J.; Group, P. W.; Sivahera, B.; Camara, M.; Kojan, R.; Walker, R.; Dighero-Kemp, B.; Cao, H.; Mukumbayi, P.; Mbala-Kingebeni, P.; Ahuka, S.; Albert, S.; Bonnett, T.; Crozier, I.; Duvenhage, M.; Proffitt, C.; Teitelbaum, M.; Moench, T.; Aboulhab, J.; Barrett, K.; Cahill, K.; Cone, K.; Eckes, R.; Hensley, L.; Herpin, B.; Higgs, E.; Ledgerwood, J.; Pierson, J.; Smolskis, M.; Sow, Y.; Tierney, J.; Sivapalasingam, S.; Holman, W.; Gettinger, N.; Vallee, D.; Nordwall, J.; Team, P. C. S., A randomized, controlled trial of Ebola virus disease therapeutics. N Engl J Med 2019, 381, 2293–2303.

6. Ashburn, T. T.; Thor, K. B., Drug repositioning: identifying and developing new uses for existing drugs. Nat Rev Drug Discov 2004, 3, 673–683.

7. Zheng, W.; Sun, W.; Simeonov, A., Drug repurposing screens and synergistic drug-combinations for infectious diseases. Br J Pharmacol 2018, 175, 181–191.

8. Pushpakom, S.; Iorio, F.; Eyers, P. A.; Escott, K. J.; Hopper, S.; Wells, A.; Doig, A.; Guilliams, T.; Latimer, J.; McNamee, C.; Norris, A.; Sanseau, P.; Cavalla, D.; Pirmohamed, M., Drug repurposing: progress, challenges and recommendations. Nat Rev Drug Discov 2019, 18, 41–58.

9. Mercorelli, B.; Palu, G.; Loregian, A., Drug repurposing for viral infectious diseases: How far are we? Trends Microbiol 2018, 26, 865–876.

10. Gil, C.; Ginex, T.; Maestro, I.; Nozal, V.; Barrado-Gil, L.; Cuesta-Geijo, M. A.; Urquiza, J.; Ramirez, D.; Alonso, C.; Campillo, N. E.; Martinez, A., COVID-19: Drug targets and potential treatments. J Med Chem 2020, 63, 12359–12386.

11. Jeon, S.; Ko, M.; Lee, J.; Choi, I.; Byun, S. Y.; Park, S.; Shum, D.; Kim, S., Identification of antiviral drug candidates against SARS-CoV-2 from FDA-approved drugs. Antimicrob Agents Chemother 2020, 64, e00819–20.

12. Riva, L.; Yuan, S.; Yin, X.; Martin-Sancho, L.; Matsunaga, N.; Pache, L.; Burgstaller-Muehlbacher, S.; De Jesus, P. D.; Teriete, P.; Hull, M. V.; Chang, M. W.; Chan, J. F.; Cao, J.; Poon, V. K.; Herbert, K. M.; Cheng, K.; Nguyen, T. H.; Rubanov, A.; Pu, Y.; Nguyen, C.; Choi, A.; Rathnasinghe, R.; Schotsaert, M.; Miorin, L.; Dejosez, M.; Zwaka, T. P.; Sit, K. Y.; Martinez-Sobrido, L.; Liu, W. C.; White, K. M.; Chapman, E.; Lendy, E. K.; Glynne, R. J.; Albrecht, R.; Ruppin, E.; Mesecar, A. D.; Johnson, J. R.; Benner, C.; Sun, R.; Schultz, P. G.; Su, A. I.; Garcia-Sastre, A.; Chatterjee, A. K.; Yuen, K. Y.; Chanda, S. K., Discovery of SARS-CoV-2 antiviral drugs through large-scale compound repurposing. Nature 2020, 586, 113–119.

13. Guy, R. K.; DiPaola, R. S.; Romanelli, F.; Dutch, R. E., Rapid repurposing of drugs for COVID-19. Science 2020, 368, 829–830.

14. Ou, X.; Liu, Y.; Lei, X.; Li, P.; Mi, D.; Ren, L.; Guo, L.; Guo, R.; Chen, T.; Hu, J.; Xiang, Z.; Mu, Z.; Chen, X.; Chen, J.; Hu, K.; Jin, Q.; Wang, J.; Qian, Z., Characterization of spike glycoprotein of SARS-CoV-2 on virus entry and its immune cross-reactivity with SARS-CoV. Nat Commun 2020, 11, 1620.

15. Walls, A. C.; Park, Y. J.; Tortorici, M. A.; Wall, A.; McGuire, A. T.; Veesler, D., Structure, function, and antigenicity of the SARS-CoV-2 spike glycoprotein. Cell 2020, 181, 281–292 e6.

16. Kam, Y. W.; Okumura, Y.; Kido, H.; Ng, L. F.; Bruzzone, R.; Altmeyer, R., Cleavage of the SARS coronavirus spike glycoprotein by airway proteases enhances virus entry into human bronchial epithelial cells in vitro. PLoS One 2009, 4, e7870.

17. Heurich, A.; Hofmann-Winkler, H.; Gierer, S.; Liepold, T.; Jahn, O.; Pohlmann, S., TMPRSS2 and ADAM17 cleave ACE2 differentially and only proteolysis by TMPRSS2 augments entry driven by the severe acute respiratory syndrome coronavirus spike protein. J Virol 2014, 88, 1293–1307.

18. Shulla, A.; Heald-Sargent, T.; Subramanya, G.; Zhao, J.; Perlman, S.; Gallagher, T., A transmembrane serine protease is linked to the severe acute respiratory syndrome coronavirus receptor and activates virus entry. J Virol 2011, 85, 873–882.

19. Dahms, S. O.; Jiao, G. S.; Than, M. E., Structural studies revealed active site distortions of human Furin by a small molecule inhibitor. ACS Chem Biol 2017, 12, 1211–1216.

20. Shang, J.; Wan, Y.; Luo, C.; Ye, G.; Geng, Q.; Auerbach, A.; Li, F., Cell entry mechanisms of SARS-CoV-2. Proc Natl Acad Sci U S A 2020, 117, 11727–11734.

21. Bosch, B. J.; Bartelink, W.; Rottier, P. J., Cathepsin L functionally cleaves the severe acute respiratory syndrome coronavirus class I fusion protein upstream of rather than adjacent to the fusion peptide. J Virol 2008, 82, 8887–8890.

22. Hardes, K.; Becker, G. L.; Lu, Y.; Dahms, S. O.; Kohler, S.; Beyer, W.; Sandvig, K.; Yamamoto, H.; Lindberg, I.; Walz, L.; von Messling, V.; Than, M. E.; Garten, W.; Steinmetzer, T., Novel Furin inhibitors with potent anti-infectious activity. ChemMedChem 2015, 10, 1218–1231.

23. Shah, P. P.; Wang, T.; Kaletsky, R. L.; Myers, M. C.; Purvis, J. E.; Jing, H.; Huryn, D. M.; Greenbaum, D. C.; Smith, A. B., 3rd; Bates, P.; Diamond, S. L., A small-molecule oxocarbazate inhibitor of human cathepsin L blocks severe acute respiratory syndrome and ebola pseudotype virus infection into human embryonic kidney 293T cells. Mol Pharmacol 2010, 78, 319–324.

24. Bekerman, E.; Neveu, G.; Shulla, A.; Brannan, J.; Pu, S. Y.; Wang, S.; Xiao, F.; Barouch-Bentov, R.; Bakken, R. R.; Mateo, R.; Govero, J.; Nagamine, C. M.; Diamond, M. S.; De Jonghe, S.; Herdewijn, P.; Dye, J. M.; Randall, G.; Einav, S., Anticancer kinase inhibitors impair intracellular viral trafficking and exert broad-spectrum antiviral effects. J Clin Invest 2017, 127, 1338–1352.

25. Stebbing, J.; Phelan, A.; Griffin, I.; Tucker, C.; Oechsle, O.; Smith, D.; Richardson, P., COVID-19: combining antiviral and anti-inflammatory treatments. Lancet Infect Dis 2020, 20, 400–402.

26. Kang, Y. L.; Chou, Y. Y.; Rothlauf, P. W.; Liu, Z.; Soh, T. K.; Cureton, D.; Case, J. B.; Chen, R. E.; Diamond, M. S.; Whelan, S. P. J.; Kirchhausen, T., Inhibition of PIKfyve kinase prevents infection by Zaire ebolavirus and SARS-CoV-2. Proc Natl Acad Sci U S A 2020, 117, 20803–20813.

27. Filippini, A.; D’Amore, A.; Palombi, F.; Carpaneto, A., Could the inhibition of endo-lysosomal two-pore channels (TPCs) by the natural flavonoid naringenin represent an option to fight SARS-CoV-2 infection? Front Microbiol 2020, 11, 970.

28. Gunaratne, G. S.; Yang, Y.; Li, F.; Walseth, T. F.; Marchant, J. S., NAADP-dependent Ca(2+) signaling regulates Middle East respiratory syndrome-coronavirus pseudovirus translocation through the endolysosomal system. Cell Calcium 2018, 75, 30–41.

29. Pruijssers, A. J.; George, A. S.; Schäfer, A.; Leist, S. R.; Gralinksi, L. E.; Dinnon, K. H., 3rd; Yount, B. L.; Agostini, M. L.; Stevens, L. J.; Chappell, J. D.; Lu, X.; Hughes, T. M.; Gully, K.; Martinez, D. R.; Brown, A. J.; Graham, R. L.; Perry, J. K.; Du Pont, V.; Pitts, J.; Ma, B.; Babusis, D.; Murakami, E.; Feng, J. Y.; Bilello, J. P.; Porter, D. P.; Cihlar, T.; Baric, R. S.; Denison, M. R.; Sheahan, T. P., Remdesivir inhibits SARS-CoV-2 in human lung cells and chimeric SARS-CoV expressing the SARS-CoV-2 RNA polymerase in mice. Cell Rep 2020, 32, 107940.

30. Weinbach, E. C.; Garbus, J., Mechanism of action of reagents that uncouple oxidative phosphorylation. Nature 1969, 221, 1016–1018.

31. Zhao, J.; He, Q.; Gong, Z.; Chen, S.; Cui, L., Niclosamide suppresses renal cell carcinoma by inhibiting Wnt/ β - catenin and inducing mitochondrial dysfunctions. Springerplus 2016, 5, 1436.

32. Agathokleous, E.; Calabrese, E. J., Hormesis: The dose response for the 21st century: The future has arrived. Toxicology 2019, 425, 152249.

33. Sun, X.; Roth, S. L.; Bialecki, M. A.; Whittaker, G. R., Internalization and fusion mechanism of vesicular stomatitis virus and related rhabdoviruses. Future Virol 2010, 5, 85–96.

34. Rasko, J. E.; Battini, J. L.; Gottschalk, R. J.; Mazo, I.; Miller, A. D., The RD114/simian type D retrovirus receptor is a neutral amino acid transporter. Proc Natl Acad Sci U S A 1999, 96, 2129–2134.

35. Chapuis, A. G.; Paolo Rizzardi, G.; D’Agostino, C.; Attinger, A.; Knabenhans, C.; Fleury, S.; Acha-Orbea, H.; Pantaleo, G., Effects of mycophenolic acid on human immunodeficiency virus infection in vitro and in vivo. Nat Med 2000, 6, 762–768.

36. Lu, S.; Pan, X.; Chen, D.; Xie, X.; Wu, Y.; Shang, W.; Jiang, X.; Sun, Y.; Fan, S.; He, J., Broad-spectrum antivirals of protoporphyrins inhibit the entry of highly pathogenic emerging viruses. Bioorg Chem 2021, 107, 104619.

37. Chan, J. F.; Kok, K. H.; Zhu, Z.; Chu, H.; To, K. K.; Yuan, S.; Yuen, K. Y., Genomic characterization of the 2019 novel human-pathogenic coronavirus isolated from a patient with atypical pneumonia after visiting Wuhan. Emerg Microbes Infect 2020, 9, 221–236.

38. Debing, Y.; Neyts, J.; Delang, L., The future of antivirals: broad-spectrum inhibitors. Curr Opin Infect Dis 2015, 28, 596–602.

39. Kumar, N.; Sharma, S.; Kumar, R.; Tripathi, B. N.; Barua, S.; Ly, H.; Rouse, B. T., Host-directed antiviral therapy. Clin Microbiol Rev 2020, 33, e00168–19.

40. Rogosnitzky, M.; Okediji, P.; Koman, I., Cepharanthine: a review of the antiviral potential of a Japanese-approved alopecia drug in COVID-19. Pharmacol Rep 2020, 72, 1509–1516.

41. Chen, C. Z.; Xu, M.; Pradhan, M.; Gorshkov, K.; Petersen, J. D.; Straus, M. R.; Zhu, W.; Shinn, P.; Guo, H.; Shen, M.; Klumpp-Thomas, C.; Michael, S. G.; Zimmerberg, J.; Zheng, W.; Whittaker, G. R., Identifying SARS-CoV-2 entry inhibitors through drug repurposing screens of SARS-S and MERS-S pseudotyped particles. ACS Pharmacol Transl Sci 2020, 3, 1165–1175.

42. Morales-Ortega, A.; Bernal-Bello, D.; Llarena-Barroso, C.; Frutos-Pérez, B.; Duarte-Millán, M.; García de Viedma-García, V.; Farfán-Sedano, A. I.; Canalejo-Castrillero, E.; Ruiz-Giardín, J. M.; Ruiz-Ruiz, J.; San Martín-López, J. V., Imatinib for COVID-19: A case report. Clin Immunol 2020, 218, 108518.

43. Weston, S.; Coleman, C. M.; Haupt, R.; Logue, J.; Matthews, K.; Li, Y.; Reyes, H. M.; Weiss, S. R.; Frieman, M. B., Broad anti-coronavirus activity of food and drug administration-approved drugs against SARS-CoV-2 in vitro and SARS-CoV in vivo. J Virol 2020, 94, e01218–20.

44. Cho, J.; Lee, Y. J.; Kim, J. H.; Kim, S. I.; Kim, S. S.; Choi, B. S.; Choi, J. H., Antiviral activity of digoxin and ouabain against SARS-CoV-2 infection and its implication for COVID-19. Sci Rep 2020, 10, 16200.

45. Cheung, Y. Y.; Chen, K. C.; Chen, H.; Seng, E. K.; Chu, J. J., Antiviral activity of lanatoside C against dengue virus infection. Antiviral Res 2014, 111, 93–99.

46. Wu, C. J.; Jan, J. T.; Chen, C. M.; Hsieh, H. P.; Hwang, D. R.; Liu, H. W.; Liu, C. Y.; Huang, H. W.; Chen, S. C.; Hong, C. F.; Lin, R. K.; Chao, Y. S.; Hsu, J. T., Inhibition of severe acute respiratory syndrome coronavirus replication by niclosamide. Antimicrob Agents Chemother 2004, 48, 2693–2696.

47. Cline, J. C.; Nelson, J. D.; Gerzon, K.; Williams, R. H.; Delong, D. C., In vitro antiviral activity of mycophenolic acid and its reversal by guanine-type compounds. Appl Microbiol 1969, 18, 14–20.

48. Schrödinger Release 2020-1; Schrödinger, LLC, New York, NY, 2020.

49. Harder, E.; Damm, W.; Maple, J.; Wu, C.; Reboul, M.; Xiang, J. Y.; Wang, L.; Lupyan, D.; Dahlgren, M. K.; Knight, J. L.; Kaus, J. W.; Cerutti, D. S.; Krilov, G.; Jorgensen, W. L.; Abel, R.; Friesner, R. A., OPLS3: A force field providing broad coverage of drug-like small molecules and proteins. J Chem Theory Comput 2016, 12, 281–296.

50. Lan, J.; Ge, J.; Yu, J.; Shan, S.; Zhou, H.; Fan, S.; Zhang, Q.; Shi, X.; Wang, Q.; Zhang, L.; Wang, X., Structure of the SARS-CoV-2 spike receptor-binding domain bound to the ACE2 receptor. Nature 2020, 581, 215–220.

51. Waterhouse, A.; Bertoni, M.; Bienert, S.; Studer, G.; Tauriello, G.; Gumienny, R.; Heer, F. T.; de Beer, T. A. P.; Rempfer, C.; Bordoli, L.; Lepore, R.; Schwede, T., SWISS-MODEL: homology modelling of protein structures and complexes. Nucleic Acids Res 2018, 46, W296–W303.

52. Case, D. A.; Ben-Shalom, I. Y.; Brozell, S. R.; Cerutti, D. S.; Cheatham, I., T. E.; Cruzeiro, V. W. D.; Darden, T. A.; Duke, R. E.; Ghoreishi, D.; Gilson, M. K.; Gohlke, H.; Goetz, A. W.; Greene, D.; Harris, R.; Homeyer, N.; Izadi, S.; Kovalenko, A.; Kurtzman, T.; Lee, T. S.; LeGrand, S.; Li, P.; Lin, C.; Liu, J.; Luchko, T.; Luo, R.; Mermelstein, D. J.; Merz, K. M.; Miao, Y.; Monard, G.; Nguyen, C.; Nguyen, H.; Omelyan, I.; Onufriev, A.; Pan, F.; Qi, R.; Roe, D. R.; Roitberg, A.; Sagui, C.; Schott-Verdugo, S.; Shen, J.; Simmerling, C. L.; Smith, J.; Salomon-Ferrer, R.; Swails, J.; Walker, R. C.; Wang, J.; Wei, H.; Wolf, R. M.; Wu, X.; Xiao, L.; York, D. M.; Kollman, P. A., AMBER 2018. 2018.

53. Maier, J. A.; Martinez, C.; Kasavajhala, K.; Wickstrom, L.; Hauser, K. E.; Simmerling, C., ff14SB: Improving the accuracy of protein side chain and backbone parameters from ff99SB. J Chem Theory Comput 2015, 11, 3696–3713.

54. Jorgensen, W. L.; Chandrasekhar, J.; Madura, J. D.; Impey, R. W.; Klein, M. L., Comparison of simple potential functions for simulating liquid water. J Chem Phys 1983, 79, 926–935.

55. Dolinsky, T. J.; Nielsen, J. E.; McCammon, J. A.; Baker, N. A., PDB2PQR: an automated pipeline for the setup of Poisson-Boltzmann electrostatics calculations. Nucleic Acids Res 2004, 32, W665–W667.

56. Ryckaert, J.-P.; Ciccotti, G.; Berendsen, H. J. C., Numerical integration fo the cartesian equations of motion of a system with constraints: molecular dynamics of n-alkanes. J Comput Phys 1977, 23, 327–341.

57. Darden, T.; York, D.; Pedersen, L., Particle mesh Ewald: An N⋅log(N) method for Ewald sums in large systems. J Chem Phys 1993, 98, 10089–11092.

58. Roe, D. R.; Cheatham, T. E., 3rd, PTRAJ and CPPTRAJ: Software for processing and analysis of molecular dynamics trajectory data. J Chem Theory Comput 2013, 9, 3084–3095.

59. Zeng, X.; Uyar, A.; Sui, D.; Donyapour, N.; Wu, D.; Dickson, A.; Hu, J., Structural insights into lethal contractural syndrome type 3 (LCCS3) caused by a missense mutation of PIP5Kgamma. Biochem J 2018, 475, 2257–2269.

60. http://research.bmh.manchester.ac.uk/bryce/amber/ (accessed March 3, 2021).

61. Meagher, K. L.; Redman, L. T.; Carlson, H. A., Development of polyphosphate parameters for use with the AMBER force field. J Comput Chem 2003, 24, 1016–1025.

62. Friesner, R. A.; Murphy, R. B.; Repasky, M. P.; Frye, L. L.; Greenwood, J. R.; Halgren, T. A.; Sanschagrin, P. C.; Mainz, D. T., Extra precision glide: docking and scoring incorporating a model of hydrophobic enclosure for protein-ligand complexes. J Med Chem 2006, 49, 6177–6196.

63. Jacobson, M. P.; Pincus, D. L.; Rapp, C. S.; Day, T. J.; Honig, B.; Shaw, D. E.; Friesner, R. A., A hierarchical approach to all-atom protein loop prediction. Proteins 2004, 55, 351–367.

64. Baik, J.; Rosania, G. R., Molecular imaging of intracellular drug-membrane aggregate formation. Mol Pharmaceutics 2011, 8, 1742–1749.

65. Gastaminza, P.; Whitten-Bauer, C.; Chisari, F. V., Unbiased probing of the entire hepatitis C virus life cycle identifies clinical compounds that target multiple aspects of the infection. Proc Natl Acad Sci U S A 2010, 107, 291–296.

66. Alley, M. C.; Scudiero, D. A.; Monks, A.; Hursey, M. L.; Czerwinski, M. J.; Fine, D. L.; Abbott, B. J.; Mayo, J. G.; Shoemaker, R. H.; Boyd, M. R., Feasibility of drug screening with panels of human tumor cell lines using a microculture tetrazolium assay. Cancer Res 1988, 48, 589–601.

67. Bartosch, B.; Dubuisson, J.; Cosset, F. L., Infectious hepatitis C virus pseudo-particles containing functional E1-E2 envelope protein complexes. J Exp Med 2003, 197, 633–642.

68. Mingorance, L.; Friesland, M.; Coto-Llerena, M.; Perez-del-Pulgar, S.; Boix, L.; Lopez-Oliva, J. M.; Bruix, J.; Forns, X.; Gastaminza, P., Selective inhibition of hepatitis C virus infection by hydroxyzine and benztropine. Antimicrob Agents Chemother 2014, 58, 3451–3460.

