## Supplemental Table and figures for "Host-directed FDA-approved drugs with antiviral activity against SARS-CoV-2 identified by hierarchical *in silico/in vitro* screening methods"

[§]Both authors contribute equally to this work

**Table S1.** Structural details about the eight targets analyzed in this study.

| Targets | Available PDB structures[a] | Selected PDB structure | Number of conformations used for VS | Binding sites | Control inhibitors |
|---|---|---|---|---|---|
| **Virus-host** | | | | | |
| **S1-RBD** | 301 | 6M0J | 6<br>- 1 Minimized<br>- 5 Clusters (0-4) from MD | **Active site:**<br>F154, Y157, N155, K85, N161, Y117, Y121, Q166, T168, N169, Y173, Y173 | **-** |
| **Host** | | | | | |
| **TMPRSS2** | - | Homology modeling from 5CE1 (TMPRSS) | 4<br>- 1 Minimized<br>- 3 Clusters (0-2) from MD | **Active site:**<br>S296, T314, D290, S291, V135, W163, C136 C152 | camostat<br>nafamostat<br>bromhexine |
| **AAK-1** | 3 | 4WSQ | 1<br>- Minimized | **Active site:**<br>G55, K74, M126, D127, F128, Q133, L183, D194 | sunitinib<br>baricitinib<br>5L4Q-ligand (**LKB**) |
| | | | 1<br>- Minimized | **Allosteric site:**<br>H82, R89, R175, D176, S197, A198, T199, N200, Q203, E208, A212, E216 | gefitinib |
| **Cathepsin-L** | 38 | 4AXL | 1<br>- Minimized | **Active site:**<br>C25, H163, N187 | oxocarbazate |
| **Furin** | 21 | 5MIM | 1<br>- Minimized | **Active site:**<br>H194, N295, S368. | 5MIM-ligand (**1n**) |
| **GAK** | 7 | 5Y7Z<br>(Active Site) | 1<br>- Minimized | **Active site:**<br>K69, E124, C126, Q129, D191 | Wee1/Chk1<br>gefitinib<br>bosutinib |
| | | 5Y80<br>(Allosteric site) | 1<br>- Minimized | **Allosteric site:**<br>D173, H200, T223, S194, N221 | gefitinib |
| **PIKfyve** | - | Homology Modeling from 6CMW (zebrafish PIP5K) | 1<br>- Minimized | **Active site:**<br>G19, K20, S21, A23, F25, I34, K36, 96, E97, N98, L99, F100, D110, K112, V132, L179, I194, D195, R198 | apilimod<br>YM201636 |
| **TPC2** | 3 | 6NQ0 (Open)<br>6NQ2 (Closed) | 2<br>- Minimized open<br>- Minimized closed | **Main site:**<br>N305, A691 | raloxifen<br>tamoxifen<br>pimozide<br>tetrandrine<br>fluphenazine<br>verapamil-S |
| | | | 2<br>- Minimized open<br>- Minimized closed | **Secondary site:**<br>W157, R210, F193 | ned19 |

[a]Last accession: October 20, 2020. Only structures for *Homo sapiens* for host-based targets were considered.

**Table S2.** Clustering analysis of MD trajectories for S1-RBD and TMPRSS2.

| | S1-RBD | | TMPRSS2 | |
|---|---|---|---|---|
| **# Clusters*** | Population | Frac | Population | Frac |
| 0 | 1721 | 43.0% | 625 | 16.1% |
| 1 | 739 | 18.5% | 606 | 15.6% |
| 2 | 595 | 14.9% | 561 | 14.4% |
| 3 | 452 | 11.3% | 406 | 10.4%** |
| 4 | 354 | 8.8% | 395 | 10.2%** |
| 5 | 62 | 1.5%*** | 367 | 9.4%** |
| 6 | 48 | 1.2%*** | 333 | 8.6%** |
| 7 | 26 | 0.6%*** | 329 | 8.5%** |
| 8 | 2 | 0.1%*** | 149 | 3.8%** |
| 9 | 1 | <0.1%*** | 115 | 3.0%** |

* A total of 10 clusters were preliminary searched. Cut-off for determining local density was 4 angstroms. Average linkage algorithm, which uses the average distance between members of two clusters, was applied.

** Discarded because the catalytic site is closed.

*** Discarded since population is less than 5%.

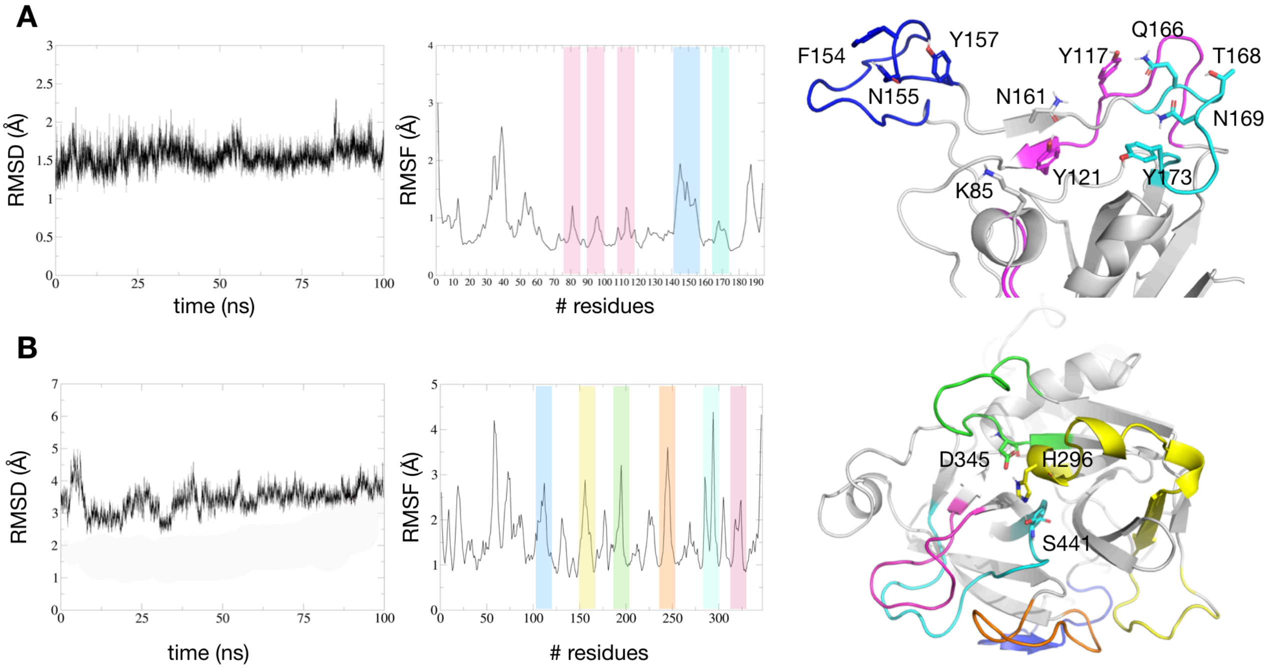


**Figure S1.** RMSD and RMSF analyses for the protein backbone atoms of (**A**) S1-RBD and (**B**) TMPRSS2. The energy minimized structure of the proteins has been used as reference for MD trajectory alignment, prior to RMSD and RMSF analyses. For both proteins, residues mainly involved in protein-ligand interactions are reported in sticks. Colored bands in the RMSF plots identify relevant regions pertaining to the binding sites.

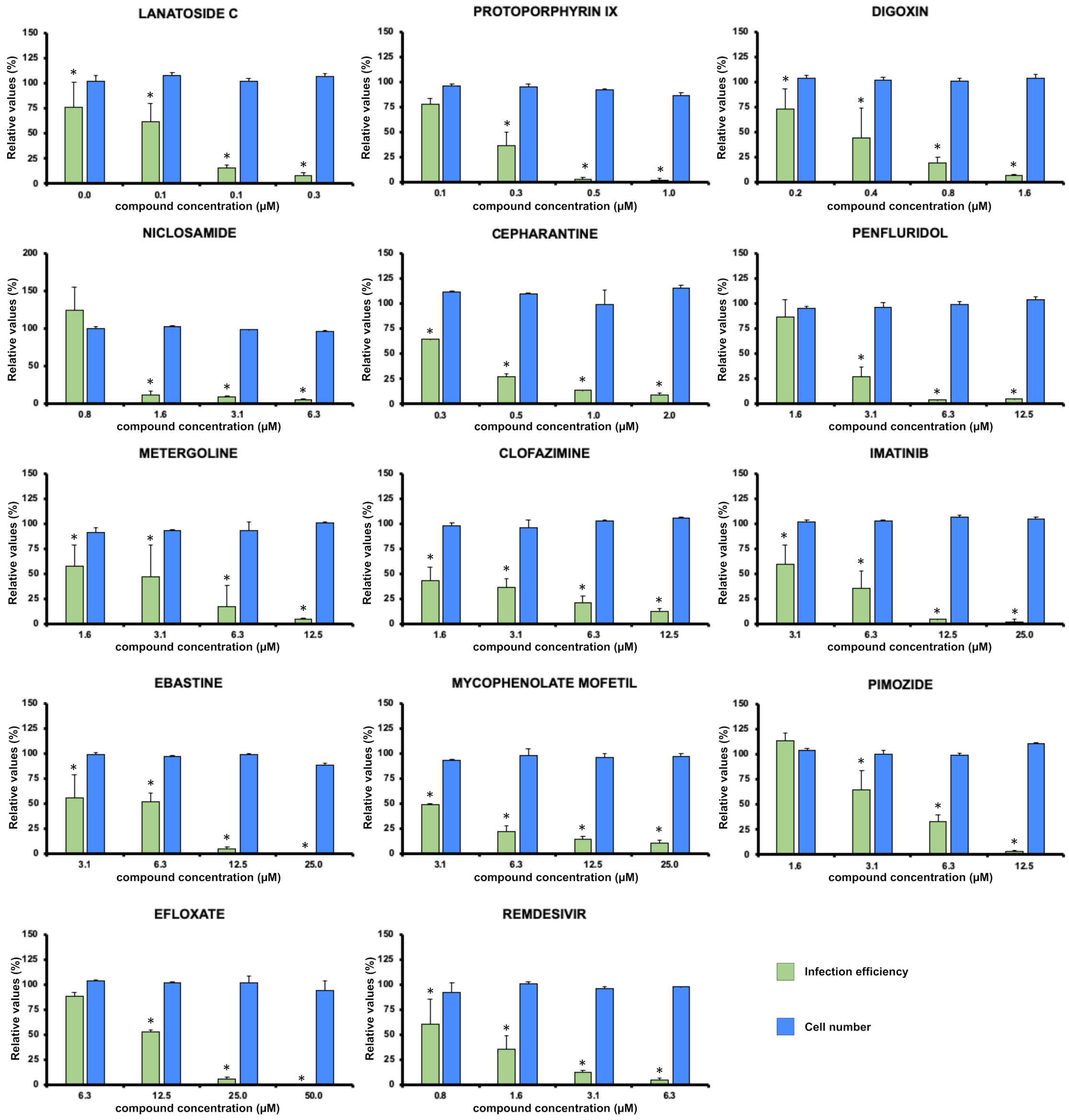


**Figure S2. Evaluation of the antiviral candidates using immunofluorescence microscopy-based viral antigen detection.** Vero-E6 cells were inoculated at MOI 0.01 in the presence of increasing compound concentrations. Relative infection efficiency was determined 24 hours post-infection by automated segmentation and signal quantitation using mock-infected cells and vehicle-treated cells as controls. Data are shown as average and standard deviation of three biological replicates. Statistical significance was determined using one-way ANOVA and Dunnet´s post-hoc test.

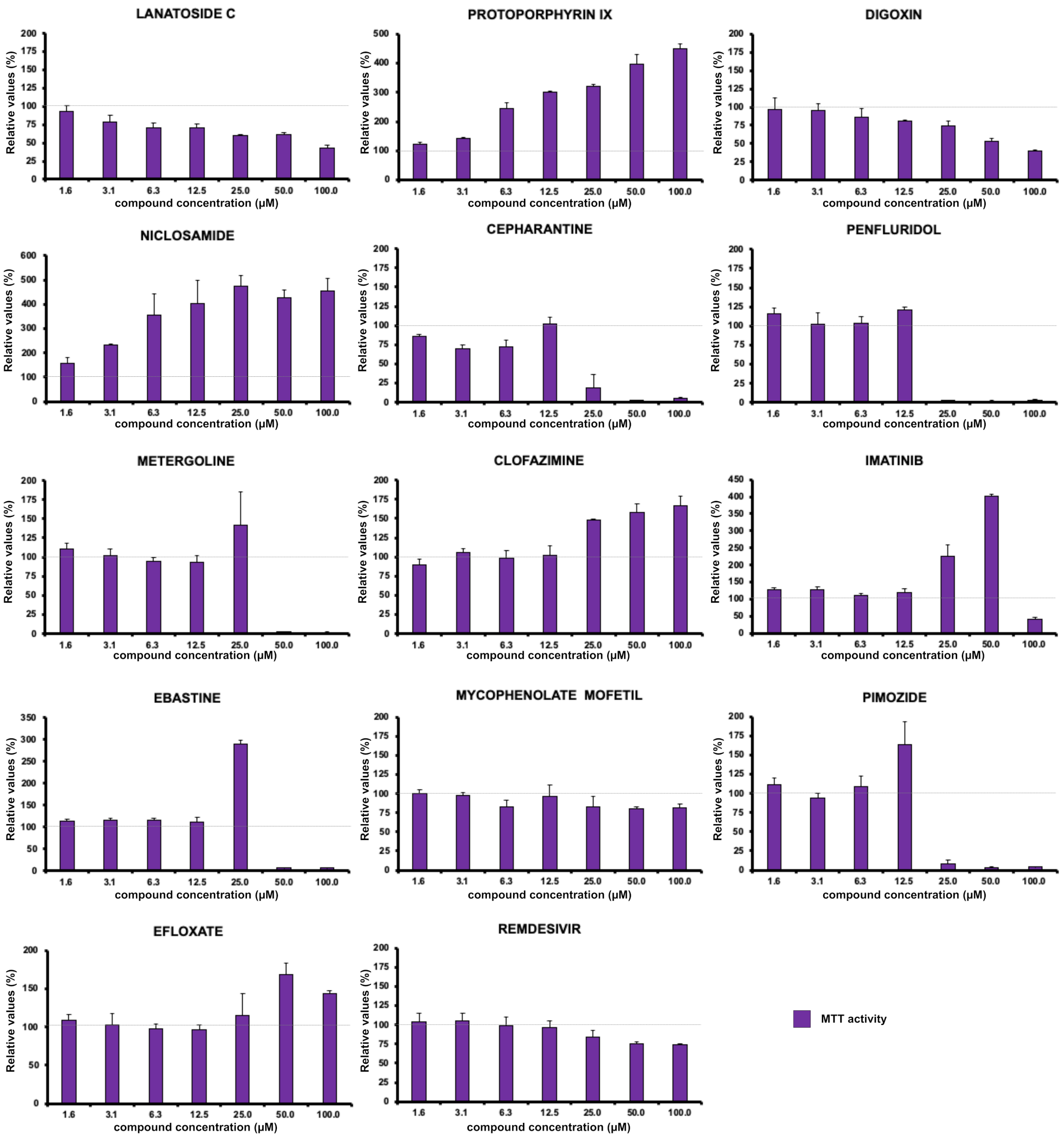


**Figure S3. Evaluation of compound cytotoxicity.** Vero-E6 cells were cultured in the presence of increasing compound concentrations. MTT activity was determined 48 hours post-treatment. Data are shown as average and standard deviation of three biological replicates.
